# Microglia perform local protein synthesis at perisynaptic and phagocytic structures

**DOI:** 10.1101/2021.01.13.426577

**Authors:** Michael J. Vasek, Jelani D. Deajon-Jackson, Yating Liu, Haley W. Crosby, Jiwon Yi, Joseph D. Dougherty

**Affiliations:** Department of Genetics, Washington University School of Medicine, 660 S. Euclid Ave, Saint Louis MO, 63108, USA; Department of Psychiatry, Washington University School of Medicine; Division of Biology and Biomedical Sciences, Washington University School of Medicine

## Abstract

Recent studies have illuminated the importance of several key signaling pathways in regulating the dynamic surveillance and phagocytic activity of microglia. Yet little is known about how these signals result in the assembly of phagolysosomal machinery near targets of phagocytosis, especially in processes distal from the microglial soma. Neurons, astrocytes, and oligodendrocytes locally regulate protein translation within distal processes. Therefore, we tested whether there is regulated local translation within peripheral microglia processes (PeMPs). We show that PeMPs contain ribosomes which engage in *de novo* protein synthesis, and these associate with a subpool of transcripts involved in pathogen defense, motility, and phagocytosis. Using a live slice preparation, we further show that acute translation blockade impairs the formation of PeMP phagocytic cups, the localization of lysosomal proteins within them, and phagocytosis. Collectively, these data argue for a regulated local translation in PeMPs and indicate a need for new translation to support dynamic microglial function.

## Introduction

Microglia serve multiple crucial functions within the central nervous system (CNS) including phagocytic “pruning” of synapses during development^1^ and disease^2,3^, clearance of apoptotic cells and debris^4^, immune surveillance, pathogen defense, and as a source of trophic factors^5^. To carry out these functions, microglia have highly dynamic peripheral microglial processes (PeMPs) which constantly extend, retract, and make brief contacts with nearby synapses, glial processes, endothelial cells, and cell somata^6–8^. During this surveillance, microglia are simultaneously sending and receiving signals through their PeMPs, yet few details are known regarding how they can align a singular nuclear transcriptional program to respond to simultaneous individualized interactions with different cellular partners at different PeMPs.

Neurons face a similar challenge: they may have thousands of synaptic inputs that need individualized responses to enable synaptic strengthening in adaptation to local activity. Thus, they utilize ribosomes present within their processes to generate proteins in response to local signals, herein referred to as “local translation.” In neurons, local translation is required for the increase in postsynaptic CamKII mediating long term potentiation^9^, and for chemically induced long term potentiation in hippocampal slices^10^. Local translation appears to be dynamically regulated by activity, influenced by untranslated region (UTR) sequences, and varied across neuronal cell types^11,12^. The regulation of neuronal local translation thus plays an important role in allowing for certain synaptic inputs to be strengthened or weakened, accordingly. Similarly, we and others have recently shown that astrocytes locally translate proteins within their distal perisynaptic and endfeet processes^13,14^. Generally, it is thought that this local translation could serve to rapidly direct specific proteins to specific spatiotemporal loci or conserve energy required to transport large proteins great distances.

Of particular interest is the rapidity by which microglia phagocytose apoptotic cells, supernumerary synapses, and pathogens during development, injury, and disease. In a live slice model, microglia engulf newly apoptotic cells within about 30 minutes^15^. During phagocytosis, actin polymerization drives rearrangement of the cytoskeleton and plasma membrane to surround the phagocytic target, forming a structure called a “phagocytic cup.” In microglia, the targets of phagocytosis may be contacted by a distal process, resulting in local phagocytic cup formation as far as 50µm from the microglial soma^15^. After engulfment of the target, the endocytosed phagosome fuses with the cellular lysosome in order to digest the contents of the phagocytized material. Interestingly, phagocytosis of zymosan particles by macrophages, which are closely related to microglia, is blocked by acute treatment with the protein synthesis inhibitor cycloheximide^16^. It is currently unknown if protein synthesis can occur locally within PeMPs, or if protein synthesis is necessary for phagocytosis by microglia.

In this study we first conducted fundamental experiments to establish whether translation occurs in PeMPs. We then developed and applied a method to profile the ribosome-bound mRNAs from PeMPs, and defined the enrichment of specific transcripts, including those involved in phagocytosis. These transcripts show enrichment of particular RNA binding protein motifs, supporting the hypothesis that local translation in PeMPs is actively regulated. Finally, we show functionally that inhibiting translation decreases the efficiency of phagocytosis, a key microglial process.

## Results

### Translation occurs in peripheral microglial processes

We first conducted a series of experiments to determine if microglia exhibit local translation within their PeMPs. If so, then we should clearly detect ribosomes and new protein synthesis within PeMPs.

To investigate whether ribosomes can be found within PeMPs, we crossed a mouse expressing the tamoxifen-inducible microglia-specific cre, CX3CR1-Cre(ERt2), with a floxed ‘TRAP’ construct mouse (ribosomal large subunit L10a fused to GFP)^17^, referred to hereafter as MG-TRAP. We then visualized the localization of ribosomes within cortical IBA1+ microglia from p28 brains. As others have reported for reporters crossed to the CX3CR1-Cre(ERt2) line^18^, all RPL10a-GFP+ cells colabeled with IBA1. Importantly for our hypothesis, we observed RPL10a-GFP signal both within the somata and the distal processes of IBA1+ cells (**Fig. 1a**). We also confirmed the presence of small ribosomal subunit RPS16 within PeMPs by immunolabeling (**Fig. 1b**). We further verified RPL10a-GFP in PeMPs through immuno electron microscopy with immuno-DAB peroxidase staining against the GFP tag, revealing ribosomal subunits within fine microglial processes located in hippocampal neuropil and apposed to neuronal presynaptic structures (**Fig. 1c**).

**Figure 1:**
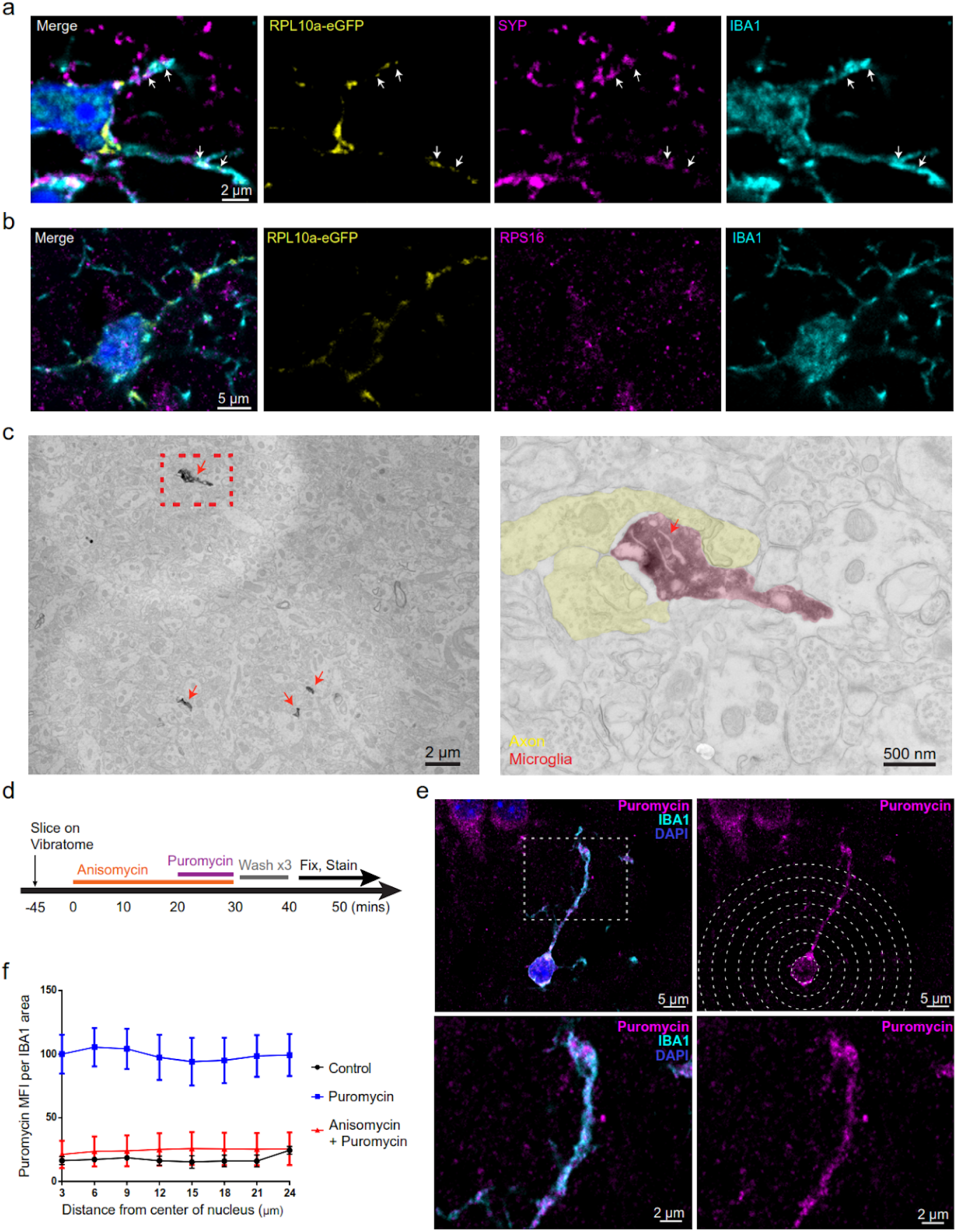
Peripheral Microglial Processes (PeMPs) contain translating ribosomes. **a-c**, Brains were harvested from 4-5 week old CX3CR1-Cre(ERT2) x RPL10a-GFP fl/fl mice, **a-b**, Immunostaining of somatosensory cortex shows localization of ribosomal subunits RPL 1Oa-GFP **(a**,**b)** and RPS16 **(b)** within distal IBA1+ processes and adjacent to synaptophysin+ **(a)** presynaptic terminals, **c**, An Immuno electron micrograph of mouse hippocampus stained with anti-GFP shows RPL10a-GFP present within fine PeMPs (pseudocolored red in zoomed inset) adjacent to several vesicle-containing axon terminals (pseudocolored yellow in zoomed inset), **d-f**, Acute live coronal slices were made from 4-5 week old C57bl6 mice and incubated as shown in the timeline **(d)**. Immunostaining **(e)** for the tRNA analog, puromycin, and IBA1 was performed to measure and quantify **(f)** mean fluorescent intensity of puromycin staining at IBA1-colocalized sites, normalized to IBA1 + area, within 3 micron radial bins from the center of the nucleus out to distal PeMPs as a readout of protein synthesis rates at increasing distances from the soma (n = 9 slices per treatment group across 3 mice processed on separate days).

Next we sought to determine if active protein synthesis occurs in these distal microglial processes. To measure and localize *de novo* peptide synthesis, we generated acute, live, 300µm slices from 4-6 week old mouse brains and incubated them with the tRNA analog puromycin, which can be used to label newly synthesized peptides^19^. As a negative control, we included slices preincubated with the protein synthesis inhibitor anisomycin, which binds to and competes for the ribosomal peptide entry site (**Fig. 1d**). These slices were then fixed and immunostained as whole mount sections. We quantified anti-IBA1 and anti-puromycin immunostaining within concentric circles of increasing diameter, centered on each microglia’s nucleus, and determined that *de novo* protein synthesis does occur in the distal processes of microglia (**Fig. 1e**). Furthermore, the mean intensity of puromycin signal (normalized to IBA1 area) was equivalent at all distances from the nucleus, as quantified in 3 µm radial bins, suggesting substantial protein synthesis occurs distal from the soma, including within PeMPs over 20 microns away. Thus, we determined that distal processes of microglia are actively engaged in *de novo* protein synthesis.

### PeMP-TRAP reveals a specific subset of microglial transcripts are enriched on peripheral microglial ribosomes

Given the presence of ribosomes and *de novo* protein synthesis, we tested whether there existed a subpool of microglial mRNA transcripts that preferentially localized to these distal, perisynaptic processes, or if all transcripts were distributed equally. We focused our studies at postnatal day 28 (p28) in order to give sufficient time for tamoxifen induction while still remaining within the period of neocortical synaptic pruning. We then isolated somatosensory cortex and hippocampus from MG-TRAP mice to generate synaptosomes using sucrose-density centrifugation^20^ in combination with translating ribosome affinity purification (TRAP) to pull down microglial ribosomes (**Fig. 2a**), following a similar protocol that has been used to identify perisynaptic neuronal^21^ and astrocytic transcripts^13^.

**Figure 2:**
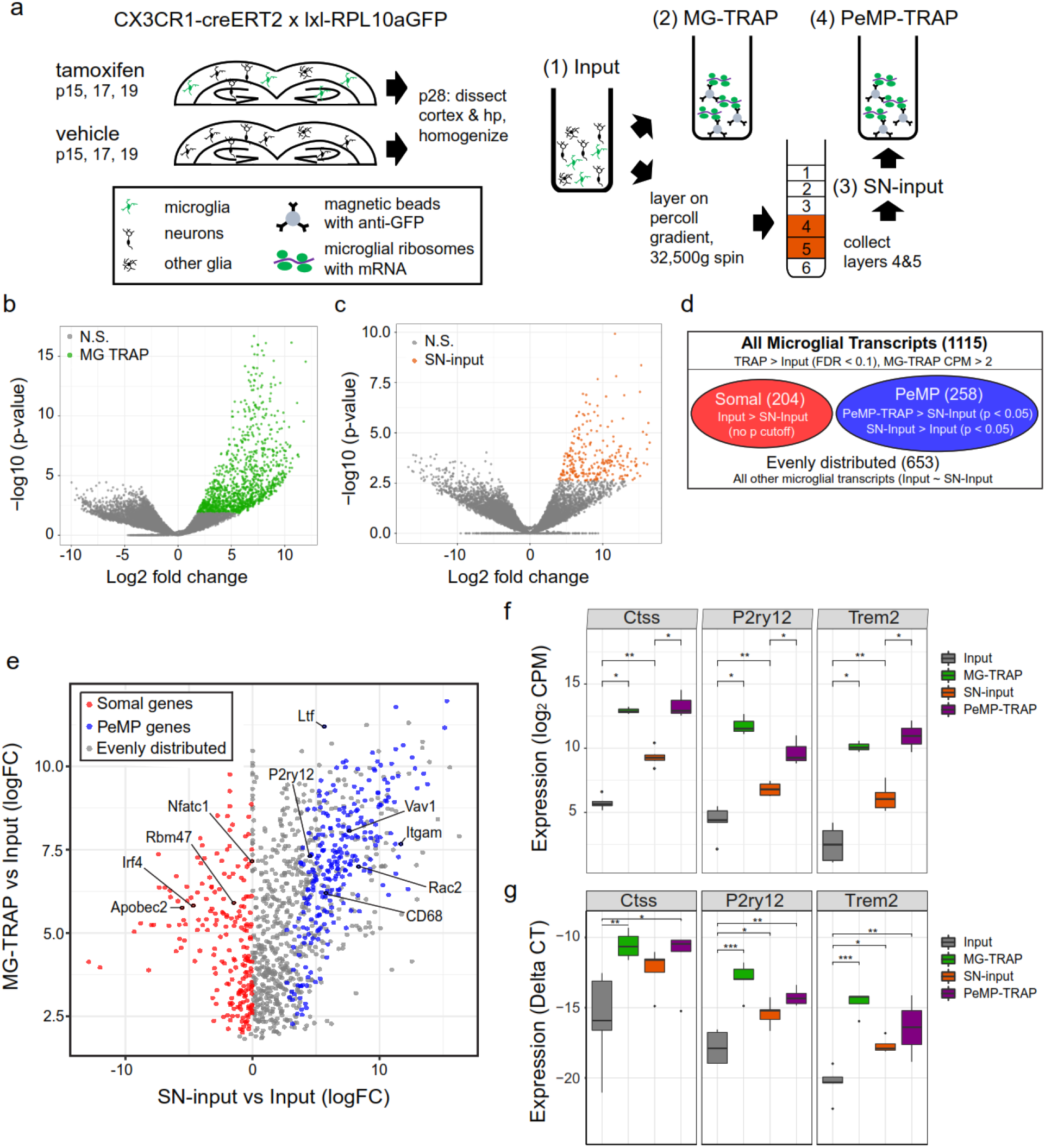
PeMP-TRAP defines transcripts enriched on peripheral microglial ribosomes. **a-g**, Dorsal cortex and hippocampus (hp) from p28 CX3CR1-CreERT2 x RPL10a-GFP(fl/fl) mice (n = 3 mixed-sex replicates of 4 pooled mouse brains each) were harvested and separated into 4 biochemical fractions per replicate, Input = dorsal cortex and hippocampal homogenate. MG-TRAP = Input subject to anti-GFP bead purification. SN-input = Sucrose density synaptosome preparation from input, PeMP-TRAP = anti-GFP bead purification of the SN-input. **b-c**, Volcano plots showing fold change and p-value of the MG-TRAP vs. Input **(b)** and SN-input vs. Input **(c)** sample comparison with genes highlighted in green or orange which passed FDR correction, **d**, Graphic depicting how PeMP and Somal microglial transcripts were defined based on comparisons between denoted biochemical fractions, **e**, Scatter plot of Log2 Fold Change (FC) in comparing the SN-input vs. Input (x-axis) and MG-TRAP vs. Input (y-axis) showing examples of several PeMP and somal microglial genes.**f**, RNAseq expression levels from each biochemical fraction of several PeMP-enriched transcripts showing enrichment in the MG-TRAP. SN-input, and PeMP-TRAP isolates, **g**, qRT-PCR was used to assess relative enrichment and depletion of perisynaptic candidates in microglial TRAP and PeMP-TRAP samples compared to input using the delta CT method normalized to 18s RNA. Statistical significance was determined by one-way ANOVA (n=5; bar\*\**p*<0.01, bar\*\*\**p<0*.*001*): CTSS: *F*=5.879, *p*=0.00664: P2ry12: *F*=9.014, *p*=0.000995; TREM2: *F*=18.62. *p*=3.7e-5; Post-hoc tests were conducted using Tukey’s HSD method (\**p*<0.05; \*\**p*<0.01; \*\*\**p*<0.001 against pre-IP).

This strategy yielded four RNA isolates per replicate for sequencing: 1) Input: whole dorsal neocortex homogenate RNA, 2) Microglial-TRAP (MG-TRAP): Anti-GFP ribosome affinity purification from the Input, 3) Synaptoneurosome-Input (SN-Input): Synaptoneurosome purification of the Input sample, 4) Peripheral Microglial Process-TRAP (PeMP-TRAP): Anti-GFP ribosome affinity purification from the SN-Input sample (**Fig. 2a**).

We first confirmed tamoxifen-dependent enrichment of microglial genes in our MG-TRAP isolates vs. the Cortex-Input isolates, as other groups have done using a similar ribotag approach (Ribosomal subunit L22-HA)^22,23^. MG-TRAP yielded a robust, tamoxifen-dependent enrichment for microglial genes among a panel of microglial, astrocyte, oligodendrocyte, endothelial, and neuronal marker genes (**Extended Data 1, 2**)^24^. Of note, MG-TRAP produced a slight enrichment for microglial genes even in the absence of tamoxifen, which corroborates other studies using the CX3CR1-Cre(ERt2) line that found a small percentage (∼2%) of IBA1+ cells exhibiting “leaky” cre reporter activity in the absence of tamoxifen^18,22^.

In comparing MG-TRAP vs Cortex-Input, we identified a list of 1115 genes significantly enriched within microglia (FDR < 0.1, MG-TRAP log2CPM > 1.0) (**Fig. 2b and Extended Data**

**3**). Our list agrees well with previously published reports of microglia transcriptomes and translatomes, encompassing all 24 microglial enriched transcripts described by Kang^23^, all 49 described by Hickman^25^, 117 of 119 described by Haimon (Odds Ratio(OR)= 717, Fisher’s Exact Test p<6.3e-126)^22^, and 342 of 486 reported by Zhang (OR=39, p< 3.3e-274)^24^.

Additionally, to confirm the SN fractionation was enriching for a separate pool of synaptic mRNA from the whole cortex we compared SN-Input to Input as done previously^12,13^. These fractions were previously found to contain fragments of a variety of cell types, including synaptoneurosomes from neurons. Consistent with this, we found that the SN-Input contained enrichment for previously characterized synaptic-localized mRNAs such as Slc1a2, Shank3, and Gsn (**Fig. 2c**). Further, a distinct subset of microglial transcripts was found in the synaptoneurosomal fraction compared to input samples, consistent with preferential localization of a subpool of microglial RNAs.

From the list of microglial transcripts (1115) we therefore designated transcripts as “PeMP-localized” if they met the following two criteria 1) PeMP-TRAP > SN-Input and 2) SN-Input > Input. As a control comparison, we designated another subset of transcripts from the microglial list as “somal” if they fulfilled the two criteria: 1) MG-TRAP > Input and 2) Input > SN-Input (**Fig. 2d, e**). The list of PeMP-localized microglial transcripts (**Extended Data 3**) includes several genes known to be involved in microglia-neuron signaling and phagocytosis such as Cathepsin S (Ctss), Trem2, and P2ry12 (**Fig. 2f)**. We then validated these findings by quantitative PCR in an independent cohort of mice (**Fig. 2g**). Analysis using Gene Ontologies (GO, **Extended Data 4**) of the PeMP-localized transcript list showed that it is enriched for transcripts involved in 122 biological processes including the immune response, cell motility, chemotaxis, phagocytosis, and synapse pruning when compared against the full MG-TRAP transcript list (**Fig. 3a**). On the other hand, the somal microglial transcript list is enriched for the biological processes of transcriptional regulation, nucleoside metabolism, and RNA modification (**Fig. 3b**) when compared against the full MG-TRAP transcript list. Examples of microglial somal-localized transcripts include Irf4, Rbm47, and the RNA-modification enzyme Apobec2 (**Fig. 2e**).

**Figure 3:**
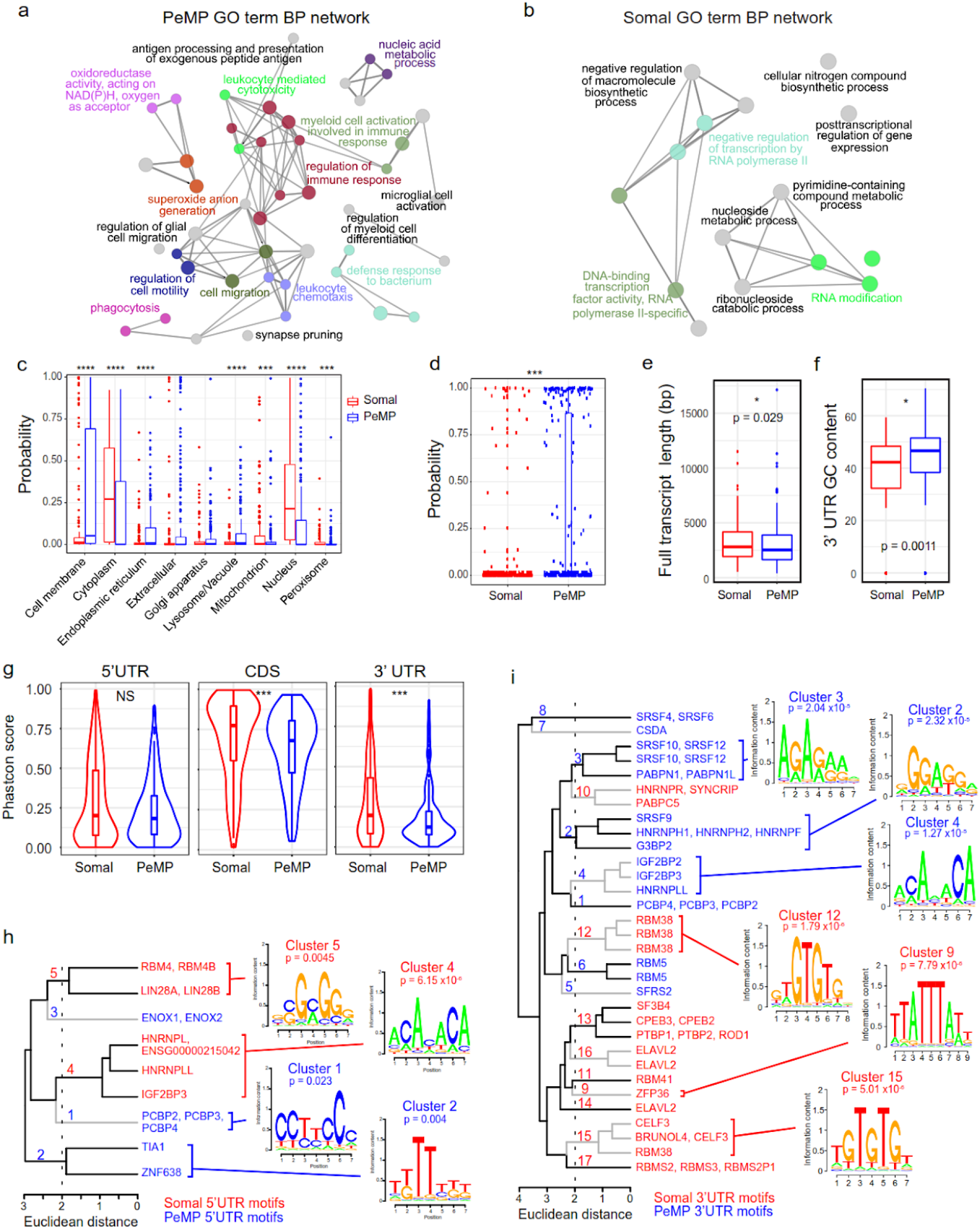
Distinguishing features of PeMP and somal enriched microglial transcripts. Network of connected and grouped Gene Ontology (GO) biological process terms generated using Cytoscape and Cluego (Bindea et al) plugin identified as enriched (**a**, p < 0.05 and **b**. p <0,02 by two-sided hypergeometric test with Benjamini-Hochberg post-hoc correction) in somal (**a**) and PeMP (**b**) candidate lists against the full microglial background list (Extended data 3), highlights distinct functions of these transcripts. Similar GO terms within the same ‘tree’ were grouped and share a color, but only the most significant GO term from each group is displayed (for full data, see Extended data 4). **c** DeepLoc analysis reveals PeMP list is enriched in transcripts predicted to localize to membranes, endoplasmic reticulum, peroxisomes, and lysosomes, while somal transcripts localize disproportionately to the nucleus, cytoplasm, and mitochondrion. (Wilcoxon ranked sum tests,*** p<0.001,****p<0.0001). **d** Predicted secretory signal peptide (Sec/SPI) are also strongly enriched in PeMP list (p = 7.6e-06, wilcoxon ranked sum test), **e** PeMP transcnpts are shorter, with higher GC content (**f**, wilcoxon ranked sum tests), and their coding sequence (CDS) and 3’UTR are less conserved by phastcon analysis (**g**, NS. not significant, *** p <0.001 by wilcoxon ranked sum test). Hierarchical clustering of motifs discovered by Transite (Krismer et al.) as significantly different in 5’ (**h**) and 3’ (**i**) UTRs illustrate average motif logos for the most significant (Fisher’s exact test, see **Extended data 5** for complete data and statistics) 2 (**h**) or 3 (**i**) clusters found in PeMP and somal lists. Cluster numbers are located on the dendrogram tree.

Local translation in other cells has been found to support subcellular localization of proteins to the correct domains^26^. Therefore, if our approach was effective in microglia, we reasoned that the PeMP-enriched transcript list should encode proteins targeted for subcellular locations near the cell’s periphery such as the plasma membrane. Conversely, the somal transcript list should encode cytoplasmic and nuclear proteins. We used the Deeploc tool^27^, which utilizes a deep learning algorithm and transcript sequence features to predict subcellular localization of transcripts, to compare the PeMP to the somal list. We found that the somal list had significantly more transcripts destined for the cytoplasm, nucleus, and mitochondrion (**Fig. 3c**), while the PeMP transcript list had significantly more transcripts destined for the cell membrane, endoplasmic reticulum, peroxisome, and of particular interest, the lysosome. Finally, we used the SignalP analysis tool^28^ and found that PeMP transcripts were more likely to encode secreted proteins based on greater signal peptide usage than somal transcripts (**Fig. 3d**).

To test the hypothesis that transcript localization may be a regulated sequence-specific process, we examined the transcripts’ 5’ untranslated region (UTR), coding, and 3’ UTR sequences to determine if there were any specific sequence features that differentiated PeMP from somal transcripts. Somal full transcripts were slightly longer than PeMP transcripts (**Fig. 3e**). PeMP transcripts had higher GC content within their 3’UTRs than somal transcripts, while GC content in the 5’UTR and coding sequence were not different (**Fig. 3f)**. Interestingly, somal transcripts’ coding and 3’UTR sequences were more highly conserved by phastcon analysis than their PeMP counterparts, but conservation of the 5’UTRs was not different (**Fig. 3g**). This suggests that some aspect of preferential localization might be due to sequence features selected to retain somal transcripts away from the periphery of the cell.

Upon discovering that certain microglial transcripts enrich on either PeMP or somal-localized ribosomes, we tested whether specific sequences underlie this spatial regulation. In neurons and many other cell types, RNA binding proteins (RBPs) bind to defined sequence motifs largely within UTRs to regulate mRNA localization, stability, and or degradation. We then utilized the Transite set motif analysis tool^29^ to test for differential RBP motif usage in the 3’ and 5’ UTRs of PeMP and somal localized microglial transcripts, focusing on motifs that showed the greatest difference in representation between those two groups (**Extended Data 5**). As motifs are often similar across RBPs, we followed this with hierarchical clustering to group similar motifs (**Fig. 3h-i**). PeMP and somal RBP motifs tended to cluster with other PeMP and somal motifs, respectively. Of particular interest, somal 3’UTRs were enriched for a TTATTTATT motif where ZFP36 (Tristetraproline) is known to bind. The ZFP36 transcript itself is also microglial- and somal-enriched, suggesting it may regulate somal transcript retention or translation. Similarly, RBM38 is another somal-encoded microglial RBP whose motif is significantly enriched within the 3’ UTRs of somal transcripts. RBP motifs significantly enriched within PeMP transcripts include the stress responsive TIA1, which is known to regulate proinflammatory genes^30,31^, as well as IGF2BP2 (also known as IMP2), a microglial and somal-enriched transcript, which is known, together with its orthologs, to regulate mRNA subcellular localization and translation during embryogenesis and early development^32,33^. While there are too many different RBPs that bind to these same motifs to motivate knockout studies at this time, the presence of such motifs strongly supports the conclusion that these gene sets contain distinct mechanisms of biological regulation.

### Protein translation blockade impedes microglial phagocytosis in ex vivo slice model

Upon discovering that transcripts encoding proteins important for phagocytosis were enriched within PeMPs, we wondered if protein synthesis was necessary for the assembly of the “phagocytic cup” (PC) structure and if key protein components of the phagolysosome might be locally synthesized rather than transported from the soma.

We first examined whether translation was indeed occurring locally within microglial PCs in an acute tissue injury model. To do so, we made fresh 300µm live brain slices from 4-5 week old mice. Near the cut surfaces of these slices, microglia display a well-characterized tissue injury response, assuming an amoeboid morphology and phagocytizing apoptotic cells and debris^15^. We briefly incubated the live slices with puromycin and measured *de novo* protein synthesis via microglial puromycin incorporation within the first 25 µm from the surface of the slice using whole-mount immunolabeling (**Fig. 4a**) as described^34^. Microglia near the slice surface developed many bulbous filopodia and structures resembling phagocytic cups at 80 minutes post-slicing (**Fig. 4b**). We found that relative puromycin levels were higher within PeMPs with phagocytic cup-like morphology (bulbous, spherical, often appearing as an IBA1+ ring) compared to PeMPs lacking these morphological features (**Fig. 4b-c**).

**Figure 4:**
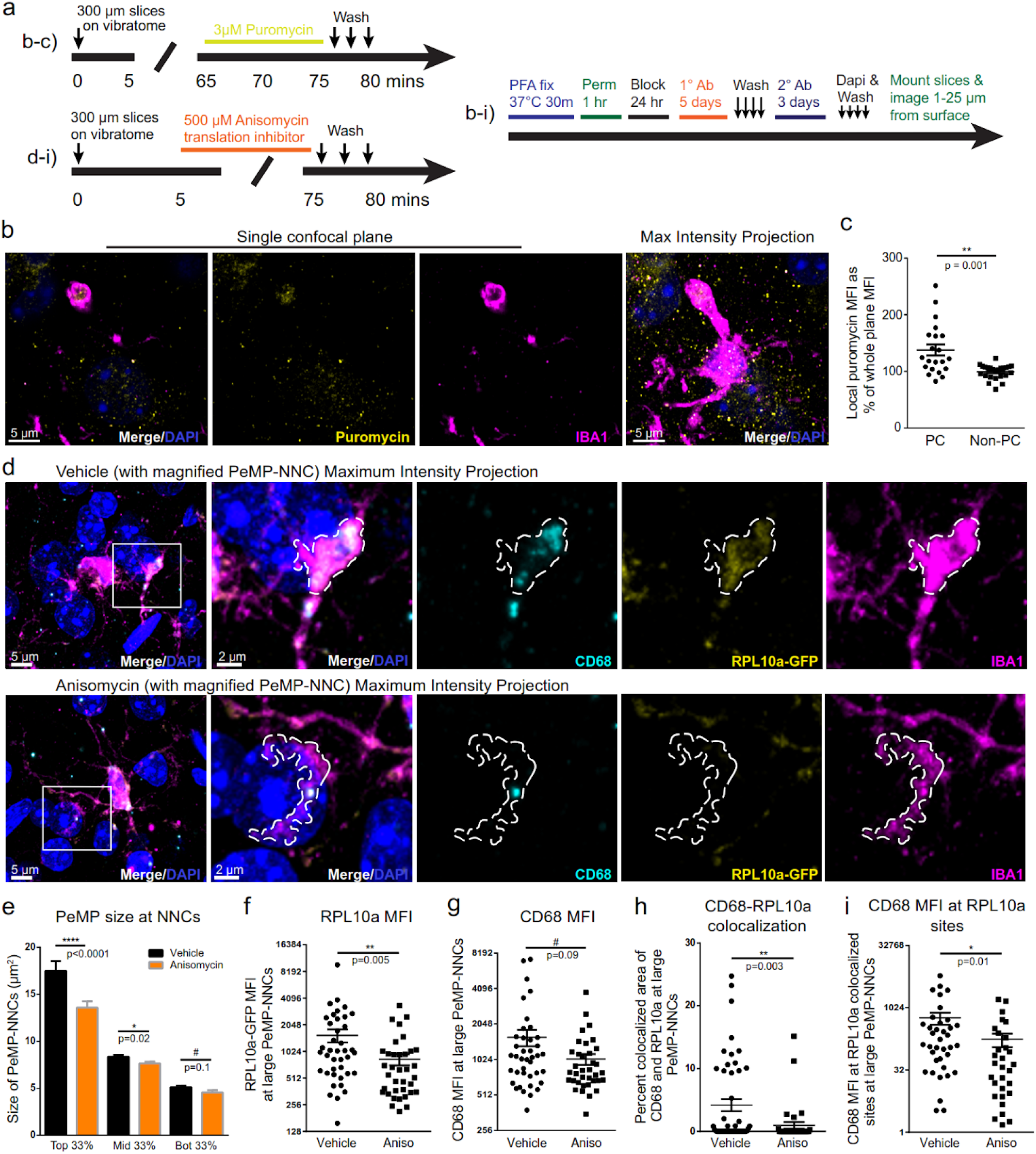
Translation is required for efficient formation of large phagocytic cups in acute slice injury model. **a**. Timelines detailing additions of puromycin or translation inhibitor, anisomycin, and tissue processing in acute live slice culture experiments (top-left applies to panels b-c, bottom-left applies to panels d-i). **b**, Whole-mount immunostain for IBA1, puromycin, and DAPI with Z-stacks taken within the first 25 μm from the surface of the slice depicts a puromycin+ phagocytic cup within a single confocal plane and as a maximum intensity projections, **c**, Quantification of local (1 μm radius) puromycin signal at puromycin-IBA1 colocalized loci, normalized to whole-plane puromycin signal, showing greater puromycin mean fluorescence intensity (MFI) at phagocytic cup (PC)-containing than non-PC containing PeMPs. ** p = 0.001 by unpaired two-tailed T-test with Welch’s correction, n = 21 and 24 PC and non-PC PeMPs, respectively from a total of 2 acute slice preps with 4 z-stacks per slice (representative of 2 independent experiments), **d**, Z-slack maximum projection images from acute-injury slices treated with vehicle or anisomycin and whole-mount immunostained for the PeMP-Iocal translation candidate, CD68. as well as RPL10a-GFP, IBA1. and DAPI taken within the first 25 μm from slice surface with area of PeMP-neighboring nuclei contacts (PeMP-NNCs) outlined in dashed line **e**, Quantification of PeMP area at PeMP-NNCs broken down by thirds (largest, middle, and smallest quantiles) per treatment group showing translation blockade impairs the formation of large area PeMP-NNCs. N = 120 (vehicle) or 109 (anisomycin) total PeMP-NNCs quantified from n = 4 independent acute slices per group and at 4 z-stacks per slice, **f-i**, Quantification of the MFI of RPL10a-GFP (f) by unpaired T-test on log-transformed data. CD68 (**g**), percent colocalized area of CD68 and RPL10a-GFP (**h**) and CD68 MFI at RPLlOa-GFP colocalized sites (**i**, unpaired T-test on log-transformed data) within largest quantile PeMP-NNCs showing dysregulation of RPL10a and CD68 localization and colocalization to large PeMP-NNCs following anisomycin translation blockade Largest quantile PeMP-NNCs consisted of N = 40 (vehicle) or 36 (anisomycin) PeMPs imaged from 4 acute slices per group (subset of n described for panel **e**) Mann-Whitney tests were used to determine significance unless otherwise noted.

To determine whether translation is required for PC formation or localization of lysosomal proteins and ribosomal subunits to PCs, we utilized MG-TRAP mouse acute live slices, blocked translation with anisomycin 5 minutes after sectioning, and then allowed the injury response to proceed for an additional 70 minutes prior to washing and fixation (**Fig. 4a**). We then performed whole-mount immunofluorescence staining for IBA1, the ribosomal subunit RPL10a-GFP, and CD68 (Macrosialin), a microglia/macrophage specific lysosomal protein and microglial local translation candidate identified via PeMP-TRAP (**Fig. 2**). We used confocal Z-stack imaging within 25µm of the slice surface and measured regions where a PeMP contacted an adjacent cell’s DAPI-stained nucleus. Per treatment group, we imaged and quantified over 100 PeMP-neighboring nuclei contacts (NNCs) connected to at least 32 microglial somata. In vehicle-treated slices, PeMP-NNCs varied in size from 2-50 µm^2^, with the larger area contacts often exhibiting cup-like three-dimensional morphologies (**Fig. 4d**) while smaller area contacts likely represent contacts without phagocytosis or where PC formation has not yet begun. PeMPs within anisomycin-treated slices also made contacts with adjacent nuclei (**Fig. 4d**), however these contacts trended slightly smaller, with very few PeMP-NNCs greater than 15 µm^2^ and more PeMP-NNCs between 0-10 µm^2^. Indeed, drug treatment specifically influenced size within the larger quantiles of contacts (**Fig. 4e**).

We then tested whether blocking protein synthesis impacted lysosomal and ribosomal protein localization within large PeMP-NNCs, measuring RPL10a-GFP and CD68. At these PeMP-nuclear contact sites, translation blockade significantly reduced levels of RPL10a-GFP (**Fig. 4f**), decreased colocalized areas of CD68 and RPL10a-GFP (**Fig. 4h**), and decreased

levels of CD68 at RPL10a-CD68 colocalized sites (**Fig. 4i**). CD68 levels, on their own, also trended lower (**Fig. 4g**). Interestingly, translation blockade did not significantly affect microglial somal MFI levels of CD68, RPL10a-GFP. or CD68 at sites of RPL10a-GFP colocalization (**Extended Data Fig. 6 a-c**). Thus, treatment with the translation inhibitor specifically decreased RPL10a and CD68 immunofluorescent signal within PeMP-NNCs.

We reasoned that loss of local CD68 signal at PeMP-NNCs during translation blockade could be due either to loss of local CD68 synthesis or to loss of a newly synthesized protein involved in transporting CD68 from the soma to the PeMP-NNC. Further, if CD68 production was occurring within the soma and then transported to processes, we would expect the somata of microglia containing many large phagocytic contacts to contain elevated levels of CD68. Therefore we traced PeMP-NNCs back to their microglial somata and measured the total area of all PeMP-NNCs belonging to each microglia as well the number of large (top third) PeMP-NNCs that each microglia manifested. We found that there was no correlation between somal CD68 levels and the total PeMP-NNC area that each microglial somata supports (**Extended Data Fig. 6d**). Furthermore, there was no significant difference in somal CD68 levels in microglia containing 0, 1, or 2 or more large PeMP-NNCs (**Extended Data Fig. 6e**). Together, these results suggest that local protein synthesis is important for microglial PC formation, consistent with local translation of CD68 at or near the PC.

Given that translation blockade in a slice injury model reduced both the size of PCs and the localization of lysosomal protein CD68 within those structures, we next tested whether microglial phagocytic function would be similarly affected. To study microglial phagocytic function, we applied phRODO-green e. Coli beads, which fluoresce strongly when internalized in an acidic cellular compartment (e.g. during phagosome fusion with the lysosome), to acute live coronal brain slices containing TdTomato+ microglia (CX3CR1-CreERT2 x AI14) (**Fig. 5a, 5b**) and live imaged ongoing phagocytosis. We found that pretreatment with the translation inhibitor, anisomycin, decreased the number of beads colocalized within microglia (**Fig. 5c**) without changing microglial area **(Fig. 5d**), consistent with impairment of phagocytosis. Taken together, these results suggest that the signaling to initiate microglial phagocytic cup formation may be translation independent, but some component of the pathway between cup formation and fusion of the phagocytic cargo with the lysosomal compartment is substantially dependent on *de novo* translation.

**Figure 5:**
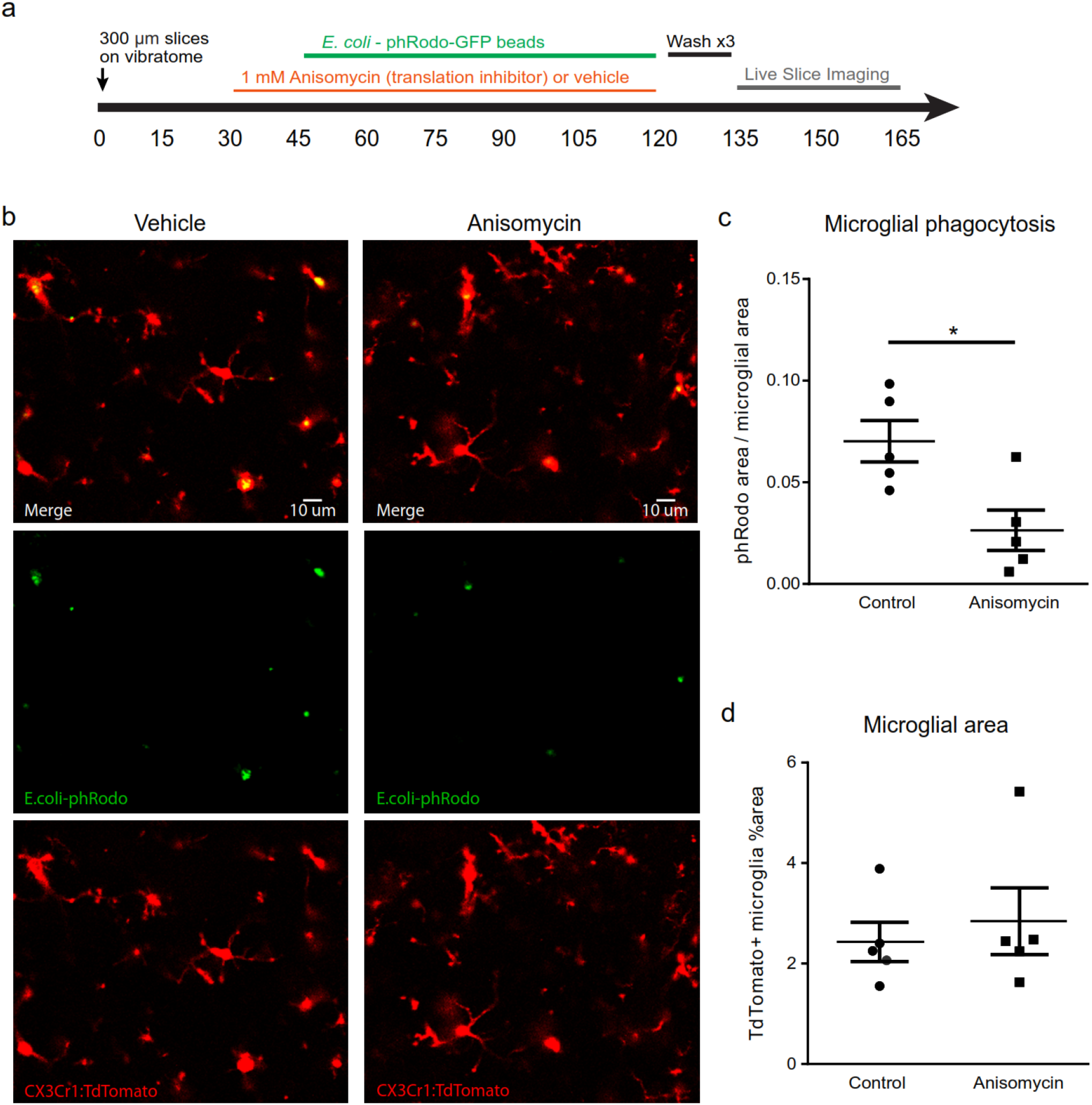
Translation is required for efficient microglial phagocytotsis. **a**, Timeline detailing experimental design, **b**, representative images from CX3CR1-CreERt2 x AI14 (floxed td Tomato) of phagocytosed phRODO-GFP E. coli beads (green), which fluoresce strongly upon acidification within the phagolysosomal compartment, within microglia (td Tomato-red) with or without the translation inhibitor anisomycin. **c**, total phagocytosis of E. coli beads, measured by phRODO-GFP and td Tomato colocalized area divided by total td Tomato area, is decreased during translation blockade with anisomycin (p = 0.015). **d**, total microglial area is unchanged (not significant, p = 0.61). N = 5 z-stacks (averaged from the 0,4, and 8 minute timepoints) taken from 2 acute brain slices per treatment group. Statistical comparisons by two-tailed unpaired t-test.

## Discussion

In this study we show that microglial processes distal from the soma contain ribosomal subunits and synthesize new proteins. These PeMP-associated ribosomes are associated with a specific subpool of microglial mRNAs which encode proteins involved in pathogen defense, phagocytosis, and cell motility. In contrast to the microglial transcripts which tend to be more restricted to the soma, the PeMP transcripts are shorter, have higher GC content within their 3’ UTRs, are less conserved, and encode more membrane, lysosomal, endoplasmic reticulum, and secreted proteins. Using an acute tissue injury model, we further show that *de novo* protein synthesis is abundant within microglial phagocytic cups (PCs). Acute translation blockade impairs PC formation, foreign particle engulfment, and the localization of ribosomal subunit RPL10a and lysosomal protein CD68 to the PC. Taken together, we describe a mechanism by which microglia may be able to translate distinct subpools of proteins in their distal processes in order to carry out specific functions, including the promotion of phagocytosis.

Transcripts enriched within PeMPs had distinct features, perhaps reflecting their distinct functions as well as hinting at mechanisms for selective enrichment on ribosomes in processes. Transcripts enriched in distal processes of neurons and astrocytes tend to be longer than their somal-restricted counterparts, so we were puzzled as to why microglial PeMP transcripts were shorter. We speculate that transcript length may be dependent upon the biological processes that these transcripts support: There is an evolutionary constraint to ensure that pathogen defense is a rapid process, especially in microglia which are key players in most CNS immune responses. Transcripts critical to this response may have evolved to be shorter in order to be transcribed, translated, and/or transported more quickly and efficiently. Likewise, the lower conservation of PeMP transcripts may reflect the enrichment for genes involved in regulation of the immune system and pathogen defense, which include many of the most polymorphic loci of mammalian genomes (e.g. MHCs, Fc Receptor genes)^35^ which have undergone rapid evolution to match the evolutionary pressure from evolving pathogens^36^. Therefore, in this case, conservation scores may not correlate with greater importance or essential function.

We report that the UTRs of PeMP and somal-enriched microglial transcripts are enriched in specific motifs, which could serve as binding sites for RBPs that either enhance or prevent their recruitment to processes. One identified motif cluster that is enriched within 3’ UTRs of PeMP transcripts includes the RBP, IGF2BP2. This family of RBPs and their homologs, including the chicken zipcode binding protein, are known to direct the localization and polarization of mRNAs within numerous cell types, especially during oogenesis and early embryogenesis^32,33^. Famously, the localization signals responsible for localizing beta-actin mRNA to the leading edge of fibroblasts^33,37^ and later to tips of neuronal growth cones^38^, were found to contain RBP motifs bound by ZBP1 and ZBP2, respectively, which are homologs to murine IGF2BP2. Here, we place IGF2BP2, the murine ZBP2 homolog, as a microglial-enriched and somal-enriched transcript whose RBP motif targets are enriched in the PeMP-enriched transcripts. This finding is consistent with a RBP that is synthesized in the cytoplasm, but performs a role in shuttling target mRNAs from the soma to the cell periphery, just as its homologs do with beta-actin. It’s also fitting that this is all taking place within PeMPs, whose motile homeostatic surveillance and phagocytic processes are both highly dependent upon actin dynamics^39^.

Prior proteomics and localization studies of purified phagosomes from macrophages, which share a common lineage with microglia, may shed some light on our findings of how *de novo*, local translation might aid phagocytosis. An electron microscopy study showed that the maturing macrophage phagosomal membrane makes contacts with endoplasmic reticulum^40^, while a 5 minute pulse-chase radiolabeling study revealed that several newly-synthesized ER-associated proteins become incorporated into the phagosomal membrane^41^. Furthermore, a mass-spectrometry study identified 19 different ribosomal subunits, including RPL10a, enriched within purified phagosomes^42^. Other proteins found to be enriched within purified phagosomes include several proteins encoded by PeMP-enriched transcripts **(Figure 2d-e, Extended data 3**) including macrosialin (CD68), macrophage capping protein (CAPG), and lysosomal hydrolases legumain (LGMN) and cathepsins D, S, and Z (CTS-D,S,Z)^42^. Importantly, expression of these PeMP-encoded hydrolases were found to peak between 1 and 24 hours post incubation with phagocytic targets^41^, consistent with a mechanism of *de novo* protein synthesis. Another PeMP-enriched transcript, myosin 1f, is known to be important for the final sealing step of the phagocytic cup around its phagocytic target^43^. Ideally, we would block translation of this transcript specifically in just one process to further test the essentiality of local translation of this protein for cup sealing, however methods to do so do not currently exist, even in neurons. Taken together, our results suggest that *de novo*, local protein synthesis near phagocytic cups is necessary for their maturation and function within microglia.

An important question raised by these studies is why microglia might utilize local protein synthesis to aid in the PeMP-enriched biological processes such as phagocytosis and pathogen defense. Since phagocytosis is a directional and location-specific process, local protein synthesis could allow a single microglia to phagocytose a nearby apoptotic cell while simultaneously contacting dozens of other healthy cells. In addition, many microglial proteins involved in these processes are expressed at low levels within healthy adult brains, but when upregulation is needed, speed is essential. Such a mechanism could be beneficial during brief windows of development, infection, or disease. In our tissue injury slice model, PC localization of CD68 and microglial phagocytosis of adjacent apoptotic cells is already visible at 80 minutes post-injury, which would be a rapid pace for transcription, capping, polyadenylation, splicing, translation, and protein transport to all occur at robust levels. One possibility is that some of these transcripts initiate translation on ribosomes and then stall, serving as a reserve of quick on-demand protein as has been demonstrated in neurons during mGluR-mediated long term depression^44^. A similar mechanism in microglia could speed up and localize debris clearance and cytokine production following injury or infection. Given that many of the PeMP transcripts produce membrane proteins, it may be informative to look for the presence of ribosome-associated vesicles in distal processes which could support this production^45^.

In the present study we found that ribosomes within peripheral microglial processes were enriched for transcripts involved in phagocytosis, and went on to show that translation blockade disrupts this process. Gene ontology analyses of transcripts loaded on PeMP ribosomes identified several additional biological processes, suggesting they may also depend on local translation, including cell motility, cell migration, immune response, and pathogen defense. It would be interesting to know if motility of distal microglial processes during homeostatic surveillance^7^ is dependent upon local protein synthesis. Furthermore, while we focused on healthy 4-5 week old mice, it would be interesting to investigate what changes occur within the PeMP-translatome using CNS disease or infection models where microglial activation, morphology, and/or phagocytosis are altered, such as Alzheimer’s Disease, Parkinson’s Disease, or the maternal immune activation model of Autism Spectrum Disorder. This approach may be able to identify transcripts that other screens would miss, whose localization or local translation at specific sites (e.g. at microglial contacts with misfolded protein deposits or synapses) are altered despite little change to total cellular levels of transcript.

## Materials and Methods

### Animals

All procedures involving animals were approved by the Washington University Institutional Animal Care and Use Committee. We utilized 4-5 week old male and female CX3CR1-Cre(ERt2) mice (official name: B6.129P2(Cg)-Cx3cr1^tm2.1(cre/ERT2)Litt^/WganJ, JAX strain 021160) crossed to cre-dependent TRAP mice (official name B6.129S4-Gt(ROSA)26Sor^tm1(CAG-EGFP/Rpl10a,-birA)Wtp^/J, JAX strain 022367) or cre-dependent tdTomato “Ai14” mice (official name: B6.Cg-Gt(ROSA)26Sor^tm14(CAG-tdTomato)Hze^/J, JAX strain 007914). Both male and female mice were used for all experiments within this study. For PeMP-TRAP RNAseq specifically, males and females were pooled in each replicate.

### Tamoxifen administration

The experimental cohort of mice was administered 100 mg/kg tamoxifen (Sigma T5648) intraperitoneally on postnatal days 15, 17, and 19, while a control cohort was administered vehicle (9:1 Sunflower oil : ethanol).

### Immunolabeling of PFA-fixed frozen brain sections

After removal, brains were stored at 4¼C in 4% Paraformaldehyde solution. After 24 hours, brains were transferred to 30% sucrose solution to soak for 48 hours, and then frozen in OCT (ThermoFisher #23-730-571) and stored at −80°C. Brains were then cryosectioned to 10 μm and applied to slides. Slides were stored at −80°C. We incubated slides with blocking solution (5% donkey serum, 0.1% Triton-X 100 in PBS) for 1 hour in a humidified chamber at room temperature. Slides were then incubated with primary antibodies overnight at 4°C in a humidified chamber. Primaries included goat anti-IBA1 (1:200), rat anti-CD68 (1:200), and mouse anti-Synaptophysin (1:50). Following this incubation, slides were washed 3x in PBS, and incubated with donkey anti-species specific alexa fluor-conjugated secondary antibodies (all at 1:400 dilution) at room temperature for one hour in blocking solution. Slides were washed 3x, then nuclei were counterstained with DAPI at 1:20,000 dilution for 5 minutes, washed once, then mounted with Prolong Gold anti-fade mounting media (ThermoFisher #P36934).

### Acute, live coronal brain slice preparation

A vibratome was used to cut 300 µm-thick coronal sections through the somatosensory cortex in ice-cold slicing buffer (125 mM NaCl, 25 mM glucose, 25 mM NaHCO_3_, 2.5 mM KCl, 1.25 mM NaH_2_PO_4_, 0.5 mM CaCl_2_, 3 mM MgCl_2_). Slices were then allowed to temporarily recover (time denoted within figures) in oxygenated, room temperature artificial cerebrospinal fluid (ACSF): 125 mM NaCl, 25 mM glucose, 25 mM NaHCO_3_, 2.5 mM KCl, 1.25 mM NaH_2_PO_4_, 0.5 Mm CaCl_2_, 3 mM MgCl_2_, and equilibrated with 95% oxygen-5% CO2.

### Acute slice treatment with puromycin and translation inhibitors

After recovery, slices were transferred to a 12-well plate with freshly oxygenated ACSF at 37°C with a gentle ∼20RPM shake. The translation inhibitor anisomycin (Sigma #A9789) was added to some slices at either 5 or 45 minutes post-slicing as noted in **figures 1d** and **4f**. To label *de novo* protein synthesis, some slices were incubated at 37°C with 3 µM puromycin (Tocris #40-895-0) at 65-75 minutes post-slicing, as noted in figures 1d and 4f.

### Live slice imaging and phagocytosis assay

Acute cortical slices (300 µm) were prepared from CX3CR1CreERT2 x AI14 (floxed TdTomato reporter) mice as described above. After recovery, slices were transferred to a 12-well plate with 1 mL freshly oxygenated ACSF and incubated with either 1mM anisomycin (Sigma #A9789) or vehicle (0.5% DMSO) at 37 °C with a gentle shake (∼20 RPM). After 15 minutes, pHrodo™ Green E. coli BioParticles™ (Thermofisher P35366, prepared according to manufacturer’s instructions and diluted in ACSF) were gently pipetted over the surface of the slices. Slices were allowed to incubate with the beads for 75 minutes at 37 °C with a gentle shake (∼20 RPM) and gently pipetting liquid from the bottom of the well over the surface of the slice every 20 minutes. Slices were then washed three times (2 minutes per wash) with 1 mL of ACSF with gentle shaking. Following washes, slices were placed in a custom perfusion chamber consisting of a silicone ring on top of a 4.5” x 6.25” sheet of no. 2 borosilicate glass (Brain Research Laboratories #5260-2) and continuously perfused with room-temperature, oxygenated ACSF while imaging. Slices were visualized on an upright Zeiss 880 confocal platform using a Zeiss 20x/1.0 water immersion objective and visible light laser excitation. Images consisting of 5-10 μm thick z-stacks at 0.76 μm step size were acquired every 2 minutes for 10 minutes using a QUASAR PMT detector.

### Immunolabeling of 300μm whole-mount coronal brain slices

Each live brain slice was washed three times (2 minutes per wash) with 1 mL of ACSF on a platform shaker, then fixed by a 30 minute incubation with 1 mL of pre-warmed 4% PFA at 37°C. Slices were then prepared with a whole-mount immunostaining procedure adapted from Dissing-Olesen and MacVicar^34^. Briefly, fixed slices (1 per well within a 12-well plate) were rinsed twice in PBS to remove residual PFA, then permeabilized using a 2-4 hour incubation in permeabilization solution (2% Triton X-100, 20% DMSO in PBS). Next, non-target epitopes were blocked through the addition of 1 mL per well of permeabilization solution with 5% donkey serum and 5% goat serum. The plate was then mixed on a shaker overnight at 60 RPM at room temperature. Each section was placed individually in a sealed custom small plastic packet containing 100 µL of a primary antibody solution with a 1:500 dilution of mouse anti-puromycin antibody (Kerafast EQ0001), 1:300 rabbit anti-IBA1 (Thermo ab5076), and 1:200 of rat anti-CD68 (eBiosciences cat#14-0681-82) in permeabilization solution with 2% donkey serum and 2% goat serum. Each packet was sealed then mixed for four days at 4°C on a 360° rotisserie wheel. Sections were then washed once per hour over 4 hours with permeabilization solution followed by a two day incubation with secondary antibody staining solution (1:400 goat anti-rabbit-Alexa-555, 1:400 goat anti-mouse IgG1-Alexa488, and 1:400 donkey anti-rat-Alexa647). All packets were wrapped with foil to prevent fluorophore exposure to light. Sections were then washed 2 additional times with permeabilization solution, stained for 20 minutes with a 1:20,000 solution of DAPI in permeabilization solution, then washed in permeabilization solution again. Immunolabeled sections were then mounted on slides as previously published^34^ except 2-3 drops of a 90% glycerol/10% PBS solution was added on top of brain slices before placing the cover slip and sealing its edges with nail polish.

### Confocal microscopy

We performed confocal microscopy on AxioImager Z2 (Zeiss). Images were taken with a 63x magnification oil lens, with an average of 1-2 microglia captured per image. Where noted in the figure legends, Z-stacks were acquired.

### Quantification of puromycin within concentric circles radiating from soma

ImageJ was used to draw concentric circles around the middle of a phagocytic cup with radii increasing by 3 µm. We created a thresholded IBA1+ cell mask and within each concentric circle we measured the mean intensity of puromycin signal at sites where it colocalized with IBA1 signal, divided by the area of IBA1+ pixels within the region.

### Quantification of puromycin, CD68 and RPL10a-GFP incorporation within microglial phagocytic structures

Following wholemount immunostaining for IBA1, CD68, and either RPL10a-GFP or puromycin, we used the polygon tool in ImageJ to demarcate the perimeter around a phagocytic cup using the IBA1 channel. The mean fluorescent intensity (MFI) of puromycin in a phagocytic cup was measured by dividing the mean puromycin signal in a cup by the fraction of the cup area containing IBA1 signal. We accounted for differences in background puromycin signal by dividing the puromycin MFI of each cup to the mean puromycin signal throughout the entire image.

### Quantification of CD68 and RPL10a-GFP incorporation at microglial-nuclei contacts

Following wholemount immunostaining for IBA1, CD68, RPL10a-GFP, and nuclear stain DAPI, we used the polygon tool in ImageJ to demarcate the perimeter of any IBA1+ process contact with a DAPI+ nucleus (excluding any contacts within 5μm of the microglial soma). The mean fluorescent intensity (MFI) of puromycin within the PeMP was measured by dividing the local mean puromycin signal by the fraction of the cup area containing IBA1 signal. We accounted for differences in background puromycin signal by dividing the local puromycin MFI of each PeMP by the puromycin MFI throughout the entire image.

### Synaptoneurosome fractionation and Translating Ribosome Affinity Purification (TRAP)

The dorsal cortex and hippocampus from p28-p29 MG-TRAP mice with or without tamoxifen administration (see tamoxifen administration) were quickly isolated and added to an ice-cold homogenization buffer as previously described^13^. All reagents and buffers used are previously described^21^. Each replicate contained the cortex and hippocampus from 4 mixed sex mice and were homogenized in 3.5 ml ice-cold homogenization buffer using a 7mL glass homogenizer (10 strokes each pestle, Kontes), and spun at 1000 × g in a Sorval RT7 for 10 min at 4°C. The supernatant from each replicate was split into two samples: For the input and MG-TRAP samples, 500 μl of the supernatant was incubated with 50 μl of Salt lysis buffer for 15 min and spun at 20,000 × g for 15 min at 4°C to reduce cell debris. The remaining supernatant (2 ml), was layered over a discontinuous 3-23% sucrose-Percoll gradient and spun at 32,500 × g for 5 min as described^20^. The synaptoneurosome band was collected by puncturing the bottom of the tube, and collecting fractions 4-5. Then, 90% of the Input and SN-Input samples (reserving 10% to purify and sequence the inputs) underwent TRAP for affinity purification of ribosomes, as described previously^12^. Briefly, samples were incubated with anti-eGFP-coated biotinylated, streptavidin MyOne T1 magnetic beads (ThermoFisher #65602) using 30 μl beads and 50 μg each of two anti-GFP antibodies (clone 19f7, 19fc8, Memorial Sloan Kettering Cancer Center antibody core) per sample. After a 4 h incubation at 4°C, samples were washed using a high salt wash and resuspended in 0.15M KCl IP wash buffer. RNA from all input and TRAP samples were then isolated and purified using a Qiagen RNEasy MinElute kit.

### Library Preparation and RNA-sequencing

RNA was assessed for quality and concentration using an Agilent RNA 6000 Pico Kit on an Agilent Bioanalyzer. RNA samples were reverse transcribed into cDNA and amplified using Smartseq v4 Ultra Low Input RNA Kit for Sequencing (Takara #634889), per the manufacturer’s instructions. The following numbers of amplification cycles (step 4) were used based on the concentration of input RNA per sample: 15 cycles for 63pg RNA input, 14 cycles for 125pg RNA input, 13 cycles for 250 pg RNA input, and 12 cycles for 500 pg RNA input. cDNA was then fragmented using a Covaris E210 sonicator using duty cycle 10, intensity 5, cycles/burst 200, time 180s, and a target insert size distribution around 200 bp was verified using an Agilent Tapestation 4200 and High Sensitivity D1000 ScreenTape and buffer (Agilent #5067-5585). Sequencing adaptors were then added on using NEBNext Ultra II DNA Library Prep Kit for Illumina (New England Biolabs #E7645S) following the manufacturer’s instructions for a 200 bp insert size distribution and using AMpure XP beads (Beckman Coulter, #A63881) for size selection and purification. Libraries were then normalized and sequenced on an Illumina HiSeq3000 as single reads extending 50 bases, by the Genome Technology Access Center at Washington University.

Sequencing results were quality checked using FastQC version 0.11.7. Illumina sequencing adaptors were removed using Trimmomatic version 0.38^46^. Reads aligned to the mouse rRNA were removed by bowtie2 version 2.3.5^47^. The remaining RNA-seq reads were then aligned to the mouse assembly (Ensembl 93) with STAR version 2.5.1a^48^. The number of uniquely mapped reads to each feature were counted using htseq-count version 0.9.1^49^. Differential expression analysis was done using edgeR version 3.32.0^50^. Only genes with CPM > 1 in at least 3 out of 24 samples were retained for further analysis. A negative binomial generalized log-linear model (GLM) was fit to the counts for each gene. Then the likelihood ratio tests (LRT) were conducted for each comparison. All code is available upon request. Raw and analyzed RNA-sequencing data are also available at the Gene Expression Omnibus (GEO), accession no.GSE161460 (reviewer token: kvshcigqbrsnryr).

### Gene Ontology (GO) term analysis and nodal networks

Following RNA sequencing, microglial somal and PeMP gene lists (Extended Data 3) were analyzed for GO term biological process (updated 06/25/2020) using the cluego plugin (v2.5.7)^51^ within the cytoscape application (v3.7.1) and using the full microglial transcript list (Extended Data 3) as the custom background. We then looked for enriched GO terms within the Somal and PeMP lists compared to the full microglial gene list via the built in two-sided hypergeometric test and Benjamini–Hochberg post-hoc correction with a cutoff of p < 0.05. A kappa score threshold of 0.4 was used to determine network connectivity. GO term tree levels included were minimum of 4 and maximum of 9. Similar GO terms were grouped with the most significant term displayed within a tree of minimum 2 common and maximum 4 different parents. Groupings are indicated by a shared color on the force directed nodal networks (**Figure 3a-b**) and in Extended Data 4. The nodal network graphic for PeMP-enriched transcripts was restricted to p < 0.02.

### RBP motif discovery using Transite transcript set motif analysis

RBP motifs were identified by entering transcript lists into the Transite transcript set motif analysis online tool (transite.mit.edu)^29^. We separately analyzed the 5’ and 3’ UTRs of PeMP and Somal gene lists using a matrix-based analysis, with the full MG-TRAP list as the background. Settings used were 50 maximum binding sites per mRNA, the transite motif database, and p-value adjustment with Benjamini-Hochberg. Enrichment scores for RBP motifs in the PeMP and Somal lists were then compared with Fisher’s exact test and false discovery rate correction to determine RBP motifs which were differentially represented within the PeMP and somal lists. The differentially represented RBP motifs were hierarchically clustered using vectorized position probability matrices (PPMs) for each motif and the hclust function with the “complete” method in R. Motif logos for each cluster were generated by seqLogo^52^ using the average PPMs across all cluster members.

### Quantitative PCR

To validate RNA transcript enrichment by PeMP-TRAP, five additional independent biological replicates of Input, TRAP, SN-input, and PeMP-TRAP were collected from p28-p35 animals as described above. cDNA samples were synthesized from RNA samples using Quanta qScript Reverse Transcriptase kit (QuantaBio, cat. no. 84002). Three technical replicates of each of the samples were quantified using iTaq Universal Sybrgreen (Bio-Rad, cat. no. 1725120) on QuantStudio 6 Flex (Applied Biosystems) in a 10 µL volume. To quantify enrichment, delta CT was calculated using 18S as an endogenous control. Data was analyzed using one-way ANOVA with Tukey’s HSD post-hoc test in R statistical software. Samples with undetermined amplification values were excluded from statistical analyses. The following primer sequences were used:

18s: F-GGGAGGTAGTGACGAAAAATAACAAT, R-TTGCCCTCCAATGGATCCT

CTSS: F-CCATTGGGATCTCTGGAAGAAAA, R-TCATGCCCACTTGGTAGGTAT (primer bank ID 10946582a1)

P2Ry12: F-ATGGATATGCCTGGTGTCAACA, R-AGCAATGGGAAGAGAACCTGG

Trem2: F-CTGGAACCGTCACCATCACTC, R-CGAAACTCGATGACTCCTCGG

### Electron microscopy

Brains from mice perfused with 4% paraformaldehyde (15710, Electron Microscopy Sciences) in PBS were excised and stored in fresh PFA fixative overnight. Coronal sections were made on a Leica VT1200S vibratome, at 100 µm to expose the hippocampus CA1 region then washed in PBS (3 × 10 min) to remove the fixative. After incubation in 0.3% hydrogen peroxide in PBS for 15 min, the sections were again washed in PBS (3 × 10 min) and placed into 1.0% sodium borohydride (NaBH4) for 15 min to quench the free aldehydes. After washing in PBS (3 × 10 min), the brain slices were incubated with agitation overnight in biotinylated anti-GFP (AB6658, Abcam) diluted 1:1000 in 2% bovine serum albumin (BSA) in PBS at 4° C. The samples were washed in PBS (3 × 10 min) the next morning and incubated for 45 min in 10ml of ABC Elite kit solution (PK-6100, Vector Laboratories) on a shaker. The slices were washed in PBS (3 × 10 min) and 0.1 M sodium acetate buffer (2 × 5 min), then incubated in nickel DAB solution (5.0 % Nickel ammonium sulfate in 0.2 M acetate buffer, 4 ml double deionized water, 1.0 ml DAB (5mg/ml) and 0.5-1.0 mg glucose oxidase) until a brown reaction product was observed (2-5 min). The brain sections were washed in PBS (3 × 10 min) and post fixed in reduced osmium (1.0% OsO4, 1.5% KCN in PBS). After washing with MQ water (3 × 10 min) the samples were stained with 2% uranyl acetate for 1 hr in the dark and washed again in MQ water (2 × 5 min) before dehydrating in a graded acetone series (50%, 70%, 90% 100% × 2 − 10 min each). Infiltration was microwave assisted (350W with vacuum) in three steps of 3 min duration (50%, 100% X2). Samples were embedded between two glass microscope slides previously treated with Teflon (MS-143XD, Miller-Stephenson) and cured at 60° C for 48 hrs. Samples were sectioned at 70nm on a Leica UC7 ultramicrotome, picked up on formvar/carbon coated grids (FCF-100-CU, Electron Microscopy Sciences), and viewed in a JEOL JEM 1400 plus TEM at 120 kV accelerating voltage.

## Acknowledgements

We would like to thank Peter Bayguinov, Gregory Strout, and the Washington University Center for Cellular Imaging for their expertise and training in live slice imaging and electron microscopy, Kristina Sakers for training and support, Claire Weichselbaum for comments on the manuscript, and the Genome Technology Access Center (GTAC@MGI) for sequencing support. Funding was provided by 5R01NS102272 and F32NS105363, and a microgrant from the Washington University Center for Cellular Imaging. GTAC is partially supported by P30 CA91842 and UL1TR002345.

## Extended Data

**Extended data figure 1.**
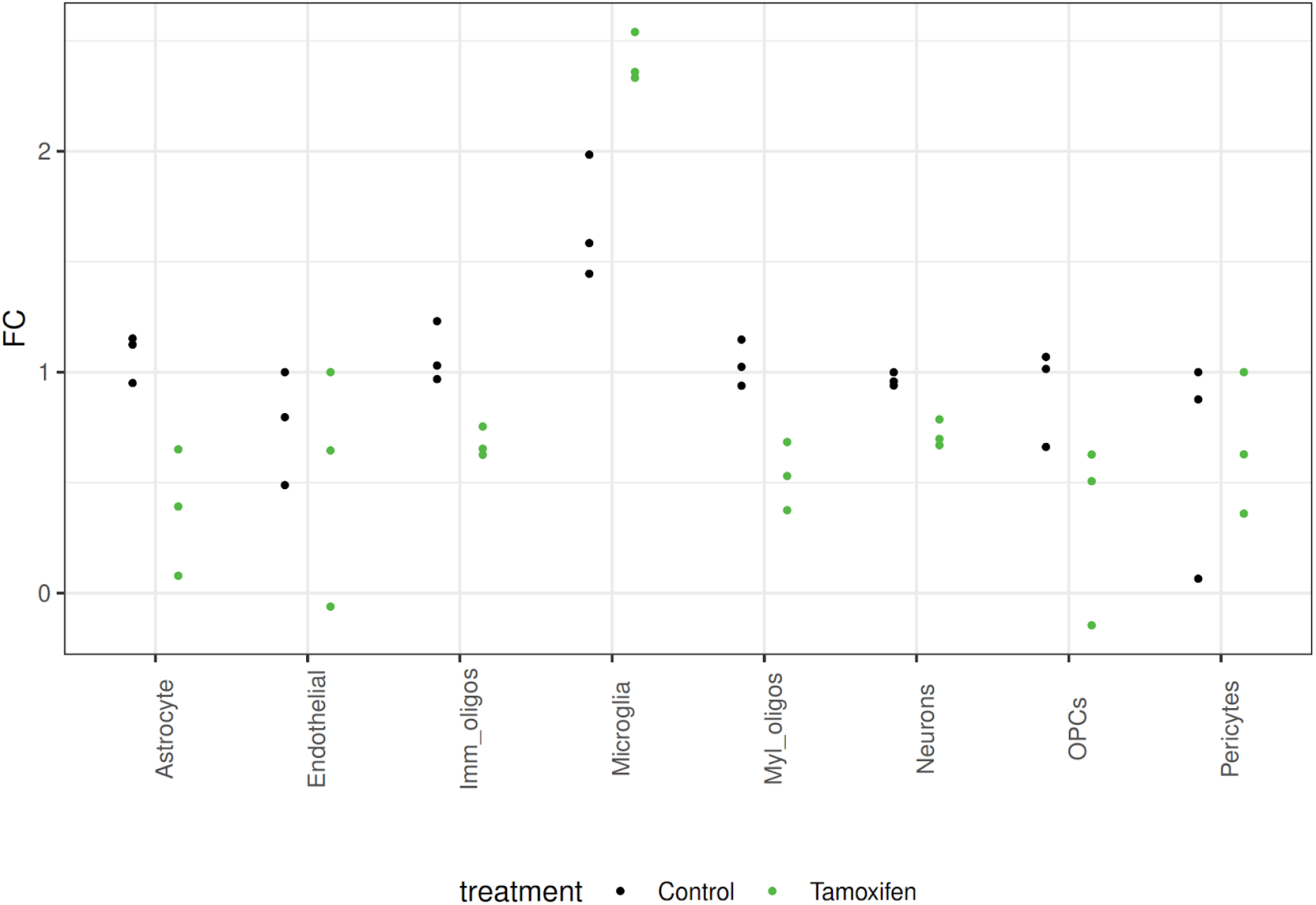
MG-TRAP yields a tamoxifen-dependent enrichment of microglial transcripts. **a)** Median fold-change expression (MG-TRAP compared to Input) of the top 20-29 marker genes per cell type (as reported by Zhang et al., see Extended Data Table 2) from replicate RNAseq experiments with (green dots) and without (black dots) tamoxifen injections reveals specific enrichment of microglial marker gene expression.

**Extended data 2. Table of CNS cell type marker genes utilized for Extended data figure 1**. Most significant 20-29 marker genes per CNS cell type generated from the web tool (https://www.brainrnaseq.org/ - compare cell type of choice vs. all other CNS cell types) described in Zhang et al.

**Extended data 3. Supplemental Table of 1115 transcripts significantly enriched by MG-TRAP, highlighting the PeMP and somal-enriched subsets**. Table contains common gene symbol, official Ensembl gene ID, and edgeR output comparing MG-TRAP to Input samples. Columns: Location: Somal, PeMP, or neither (NA) as defined in methods. logFC: log (base 2) fold change for enrichment in MG-TRAP compared to input. logCPM: log counts per million LR: Likelihood ratio test, with corresponding p-value and FDR (Benjamini-Hochberg) correction. Remaining columns are mean log (base 2) CPM for individual sample types. Only samples with log_2_ CPM > 1, FC > 0, and corrected p-value <0.1 were included. Unfiltered data is available at the Gene Expression Omnibus (GEO), accession no.GSE161460 (reviewer token: kvshcigqbrsnryr).

**Extended data 4. Supplemental Table of clueGO full output for GO term biological process PeMP vs MG-TRAP and Somal vs MG-TRAP Genes**. GO term biological processes associated with PeMP and somal-enriched (separate tabs) microglial genes was calculated using the ClueGO plugin (Bindea et al 2009) for Cytoscape (See methods for settings) using the full microglial gene list (**Extended data 3**) as a custom background Column headers are as defined by clueGO. Only GO terms with Benjamini-Hochberg corrected p < 0.05 are listed. Coloration/term groupings match Figure 3a-b. Within the PeMP-enriched list, GO term grouping was only carried out on terms with a corrected p value < 0.02 in order to match Figure 3a.

**Extended data 5. Supplemental Table of nucleotide motifs enriched in Somal and PeMP gene lists**. Column headers are as defined by Transite output(Krismer et al., 2020). Transite was run comparing PeMP list to all TRAP enriched genes (columns D-F, J), and Somal list to all TRAP enriched genes (columns G-H, K). A Fisher s Exact test (column K) was used to identify motifs found more in PeMP or Somal lists than expected by chance, and corrected for multiple testing by Benjamin-Hochberg FDR (column P). Only FDR< 05 were retained. Ratio of motif enrichments was also calculated (column M). Results of hierarchical clustering (based on motif sequences Position Weight Matrices), are also included (columns R, S).

**Extended data 6.**
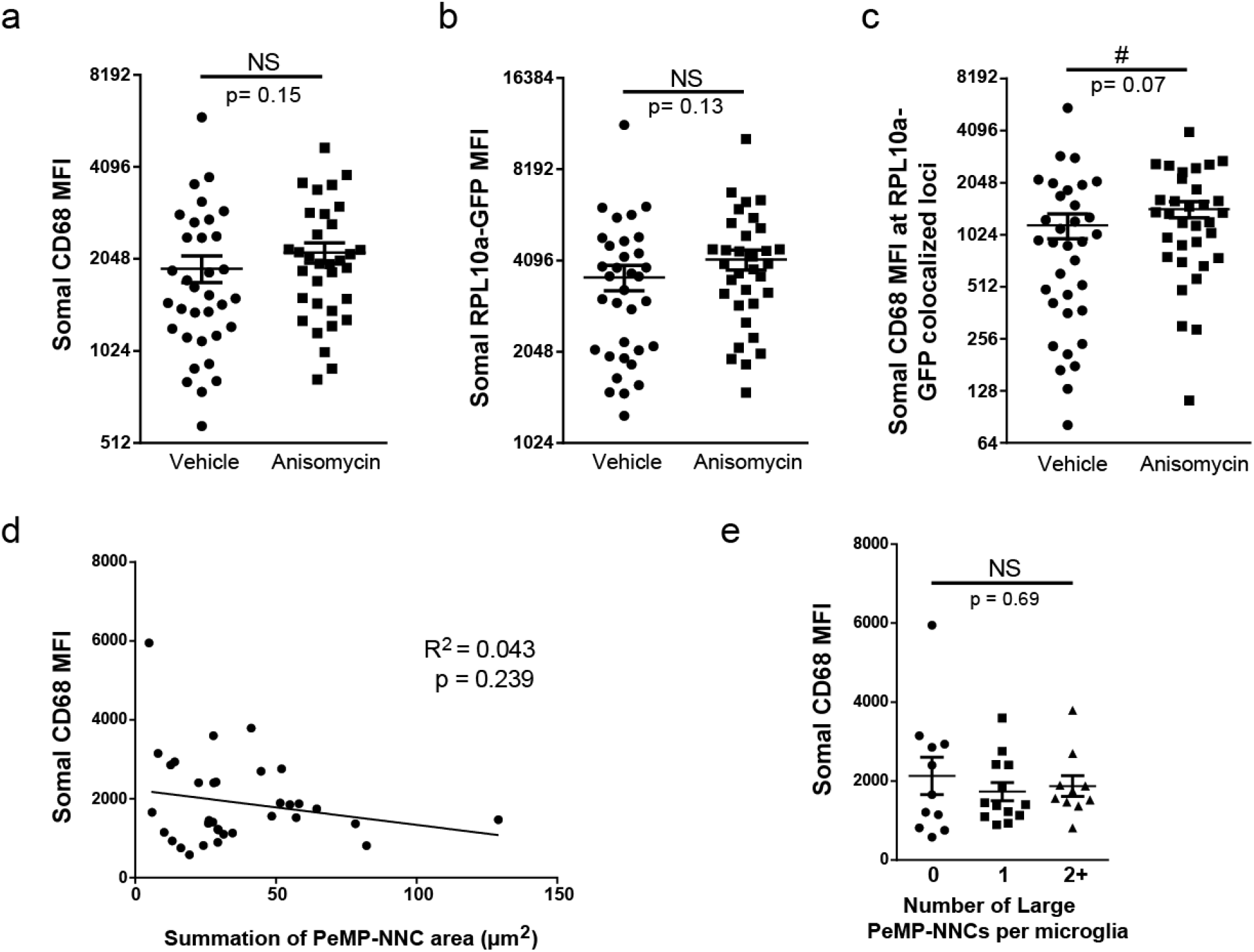
Translation blockade or number of contacts does not affect somal CD68 or RPL10a expression. **a-e)** Whole mount immunostains were prepared from acute slices as in Figure 4d-i. **a-c**, No significant differences were detected between anisomycin and vehicle-treated slices in somal mean fluorescence intensity (MFI) levels of CD68 (**a**. p = 0.15 by unpaired two-tailed t-test on Iog2 transformed data), RPL lOa-GFP (**b**, p = 0.13 by unpaired two-tailed t-test on log2 transformed data), and CD68 at RPl10a-GFP colocalized loci (**c, # p** = 0.07 by Mann-Whitney) **d**, There was no correlation between vehicle-treated microglial somal CD68 MFI and the summation of its PeMP-NNC areas. R^2^ = 0.04 and slope not-significantly different from zero, p = 0.24 by F-test. **e**, There was no statistical difference between vehicle-treated microglial somal CD68 MFI among those supporting either 0, 1. or 2 or more large (top third quantile) PeMP-NNCs. p = 0.69 by one-way AN OVA. **a-e**, N = 34 (vehicle) or 33 (anisomycin) microglial somata imaged from 4 independent slices per group with at least 4 z-stack images per slice.

